# Anisotropic light propagation in human brain white matter

**DOI:** 10.1101/2025.04.02.646745

**Authors:** Ernesto Pini, Danila Di Meo, Irene Costantini, Michele Sorelli, Samuel Bradley, Diederik S. Wiersma, Francesco S. Pavone, Lorenzo Pattelli

**Affiliations:** Istituto Nazionale di Ricerca Metrologica (INRiM), 10135, Turin, Italy; European Laboratory for Non-linear Spectroscopy (LENS), 50019, Sesto Fiorentino, Italy; Department of Biology, University of Florence, Via Madonna del Piano, 6, Sesto Fiorentino 50019, Italy; Department of Physics and Astronomy, Universitàdi Firenze, 50019, Sesto Fiorentino, Italy; National Research Council – National Institute of Optics (CNR-INO), 50019, Sesto Fiorentino, Italy

**Keywords:** multiple scattering, time-resolved measurements, anisotropic diffusion, scattering tensor, neurophotonics

## Abstract

**Significance:** Accurate modeling of light diffusion in the human brain is crucial for applications in optogenetics and spectroscopy diagnostic techniques. White matter tissue is composed of myelinated axon bundles, suggesting the occurrence of enhanced light diffusion along their local orientation direction, which however has never been characterized experimentally. Existing diffuse optics models assume isotropic properties, limiting their accuracy.

**Aim:** We aim to characterize the anisotropic scattering properties of human white matter tissue by directly measuring its tensor scattering components along different directions, and to correlate them with the local axon fiber orientation.

**Approach:** Using a time- and space-resolved setup, we image the transverse propagation of diffusely reflected light across two perpendicular directions in a ex vivo human brain sample. Local fiber orientation is independently determined using light sheet fluorescence microscopy (LSFM).

**Results:** The directional dependence of light propagation in organized myelinated axon bundles is characterized via Monte Carlo (MC) simulations accounting for a tensor scattering coefficient, revealing a lower scattering rate parallel to the fiber orientation. The effects of white matter anisotropy are further assessed by simulating a typical time-domain near-infrared spectroscopy measurement in a four-layer human head model.

**Conclusions:** This study provides a first characterization of the anisotropic scattering properties in ex vivo human white matter, highlighting its direct correlation with axon fiber orientation, and opening to the realization of quantitatively accurate anisotropy-aware human head 3D meshes for diffuse optics applications.

## 1 Introduction

The intricate macroscopic and microscopic structure of the human brain poses a significant challenge for techniques that rely on diffuse near-infrared light for functional imaging and diagnostics.^1^ The complex and heterogeneous composition of brain tissue influences light propagation, affecting the accuracy of these techniques. Current optical methods, such as functional near-infrared spectroscopy (fNIRS) and diffuse optical tomography (DOT), focus primarily on monitoring hemodynamic responses by mapping blood oxygenation changes in different brain regions.^2–4^ Among these, time-resolved techniques play a crucial role in extracting absolute tissue oxygen saturation values and estimating the reduced scattering coefficient,^5^ both of which are essential for functional and clinical assessments. A key advantage of these methods is their ability to separate extra-cerebral and cerebral contributions, as different photon propagation times correspond to different penetration depths. This feature enhances the accuracy of functional imaging and disease diagnostics by isolating signals from deeper brain structures.

In addition to neuroimaging, light diffusion in brain tissue is also relevant to optogenetics, a technique enabling precise, cell-specific control of neural circuits using light-sensitive proteins. Optogenetics allows to stimulate or inhibit neurons with high temporal and spatial precision,^6, 7^ which however requires accurate modeling of light transport in neural tissue to ensure that light delivery reaches the desired depth and intensity for activating targeted cells. The high scattering of brain tissue significantly influences the illumination profile, affecting the efficiency of optogenetic activation, especially for deep-brain stimulation via optical fibers. While previous studies examined how light propagates through brain tissue,^8^ isotropic properties are often assumed in spite of the apparent structural anisotropy of white matter tracts.

White matter is composed of myelinated axons that are locally organized to form bundles. The preferential alignment of these bundles creates a directional dependence in light diffusion, as elongated structures interact differently with light depending on its propagation direction. This anisotropic behavior has been well-characterized in diffusion-weighted magnetic resonance imaging (DW-MRI) and diffusion tensor imaging (DTI), where axonal structures are mapped based on water diffusion patterns.^9^ However, anisotropy in diffuse optics remains largely disregarded, despite its potential impact on optical neuroimaging and optogenetics, with a lack of experimental and numerical validation for anisotropic scattering coefficients in human white matter.

A major challenge in modeling brain tissue optics is the scarcity of widely accepted reference values for optical properties, which complicates the development of standardized models. Experimental studies report highly variable results,^10–15^ often due to differences in sample preparation (e.g., formalin fixation, paraffin embedding)^16^ or measurement conditions such as temperature-dependent scattering changes.^17, 18^

In this scenario, studies in the field of diffuse optics acknowledging the structural anisotropy of white matter are scarce. Hebeda et al.^19^ reported a first measurement of the effective attenuation coefficient (defined as 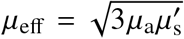) along parallel and perpendicular directions with respect to white matter axons in bovine brain, finding significant directional differences: *μ*_eff,∥_ = (0.47 ± 0.06) mm^−1^, *μ*_eff,⊥_ = (0.63 ± 0.13) mm^−1^ at 633 nm. Heiskala et al.^20,21^ attempted a description of anisotropic light transport in the whole human head using a simplistic version of the anisotropic diffusive equation and MC simulations using parameters derived from DTI measurements. While this approach allowed to recognize the presence of transport anisotropy in white matter, it failed to accurately predict its effects in single frequency-domain diffuse optical tomography measurements. Additionally, no attempt was made at measuring the actual direction-dependent scattering coefficients, which were simply assumed proportional to DTI fractional anisotropy data for water diffusion through white matter tissue. More recently, De Paoli et al.^22^ measured direction-dependent reduced scattering coefficients in spinal cord white matter, reporting a large transport anisotropy associated with the high degree of alignment of axons in the spinal cord. Relevant works in this direction have also been published by Menzel et al.,^23, 24^ who modeled the anisotropic geometry of individual nerve fibers bundles using Finite-Difference Time-Domain simulations, and showed that the angular distribution of light scattered from ultrathin brain slices depends on the local axon orientation. These results are also of high interest to understand whether myelinated axon fibers can favor the directional propagation of biophotons in brain tissue, a topic which received considerable attention in recent years.^25, 26^

Despite these efforts, accurate experimental characterization of the direction-dependent scattering coefficient inside human brain white matter is still lacking. In this work, we introduce a tensor-based model for the description of anisotropic light transport, enabling us to overcome previous oversimplifications of scattering in structurally anisotropic media. Our experimental characterization spans time scales covering both diffusive propagation regimes and sub-diffusive transients, providing new insights into the optical properties of white matter, with implications for neuroimaging, optogenetics and computational light transport models.

## 2 Anisotropic diffusion model

In anisotropic materials the diffusive constant *D* and the scattering coefficient *μ*_s_ can be represented as 3 × 3 tensor quantities ***D, μ***_s_ which can be usually diagonalized using an appropriate common reference frame. In analogy to the isotropic case, the diffusion tensor elements can be associated to reduced scattering coefficients along different directions 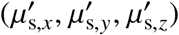:

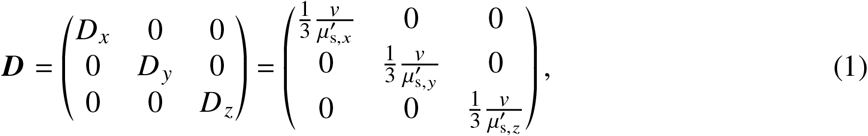

having dimensions of [m^2^ s^−1^], where *v* = *c*/*n*_eff_ represents the speed of light in the medium, with *c* as the speed of light in vacuum and *n*_eff_ as the medium effective refractive index.

On the other hand, at the microscopic level, we can define a diagonal scattering coefficient tensor

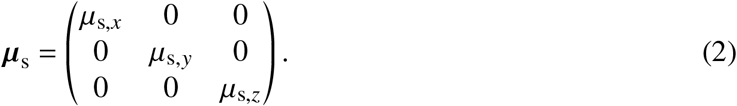

In the diffusive regime, analytical relations between these two tensors have been recently derived for the case of uniaxial structural asymmetry^27^ typical of aligned fiber bundles, revealing a non-trivial dependence of diffusion rates along each individual axis, from the microscopic scattering coefficients in all three directions. Compared to the simplistic assumption that each component of the diffusion tensor depends exclusively on its corresponding scattering coefficient, the exact solution allows to avoid systematic errors, the magnitude of which is directly related to the degree of structural anisotropy.^28^

In general, before the onset of diffusion, scattering asymmetry is also expected to influence light transport, as often modeled through the Heyney-Greenstein phase function with an asymmetry factor *g*. In analogy with ***μ*** and ***D***, the asymmetry factor itself could take a tensor form in anisotropic media, which however would double the number of free parameters and make the quantitative retrieval of optical properties unfeasible. In diffusive isotropic media, this is typically not a problem thanks to the existence of a simple “similarity relation” linking the scattering coefficient to its reduced counterpart 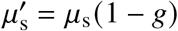. However, this simple relation does not hold in the anisotropic case, and the microscopic interpretation of the reduced scattering coefficient in the random walk picture of diffusion is lost. To restore a microscopic quantity of practical utility, we introduce a trade-off by considering a tensor scattering coefficient and a scalar asymmetry factor, which we combine into an *effective reduced scattering coefficient* 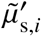:^29, 30^

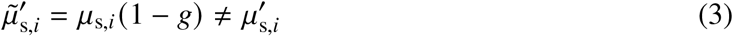

with *i* = {*x, y, z*}. The effective reduced scattering coefficient accounts for the directional dependence of scattering in anisotropic media, providing a more accurate description of light transport especially in the case of biological tissues, where the asymmetry factor is expected to be close to unity irrespective of the direction.

The main aim of this study is to quantitatively determine the components of the 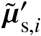 tensor and of the *g* scalar by looking at light transport dynamics during the early transient and the subsequent diffusive regime.

To quantify the degree of light diffusion anisotropy, we further introduce an *Optical Fractional Anisotropy* (*OFA*), inspired by the concept of fractional anisotropy (*FA*) commonly used in the field of diffusion tensor imaging^31^ (based on water water molecule diffusion) or in Structure Tensor (ST) analysis^32^ (based on histological sections microscopy). The Optical Fractional Anisotropy follows the same definition, using the components of the light diffusion tensor:

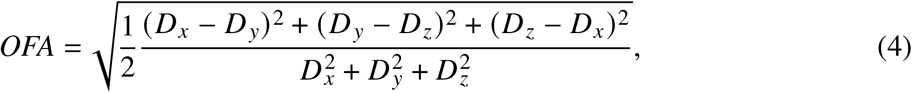

and is similarly comprised between 0 and 1 for isotropic diffusion and perfectly directional propagation, respectively.

## 3 Materials and Methods

### 3.1 Sample preparation

Human tissue sample were procured through the body donation program (Association des dons du corps) of the Université de Tours. Written consent was obtained from healthy participants prior to death, including the brain for any educational or research purposes. The authorization documents of the Association des dons du corps are kept with the Body Donation Program at the Université de Tours, and were collected within the general frame of the approved IRB submission to the Partners Institutional Biosafety Committee (PIBC., protocol 2003P001937). Upon collection, samples were placed in neutral buffered formalin (pH 7.2–7.4) and stored at room temperature. Blocks from the fixed samples were washed with phosphate-buffered saline solution at 4 °C with gentle shaking for 1 month.

### 3.2 Optical gating setup

Our study considered diffuse reflectance from a piece of post-mortem human brain pericalcarine cortex with dimensions of roughly 18 × 15 × 20 mm, presenting both gray and white matter regions (Fig. 1a). Utilizing near-infrared light at *λ* = 820 nm, we performed time- and space-resolved reflection measurements on the white matter region due to its known fibrous structure composed of locally aligned bundles of neuron axons. Anisotropic light transport was studied by means of a transient imaging apparatus based on an optical gating scheme (Fig. 2) allowing to capture the spatio-temporal evolution of reflected intensity with sub-ps resolution^30, 33^ upon illumination with a focused pulse with a duration of ∼150 fs. The laser sources emit trains of synchronous pump-gate pulses at different wavelengths, which are recombined before detection in a *β*-barium borate (BBO) non-linear crystal, where a sum-frequency conversion occurs producing a signal at a visible wavelength. The gate beam is expanded to achieve a flat intensity distribution over the whole non-linear crystal surface to obtain a uniform upconversion efficiency of the signal over the whole field of view. A motorized delay stage is used to adjust the relative delay between the probe and gate arms of the setup, thus allowing to resolve the fast dynamics of light diffusion.

**Fig 1:**
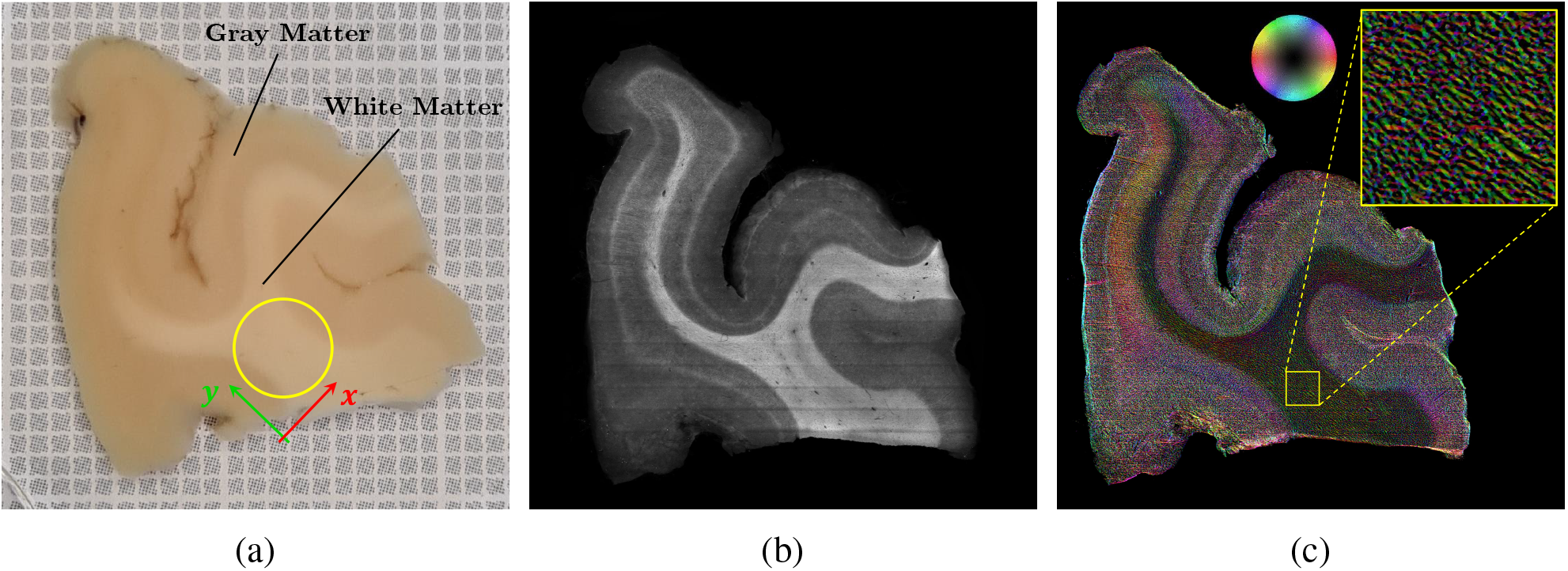
(a) 1 mm-thick brain slice placed on millimeter grid paper for reference. The yellow circle represents the circular field of view selected for subsequent measurements. (b) Raw acquisition image obtained with light-sheet fluorescence microscopy after clearing. (c) Image of the axon fiber orientation 3D map. The color wheel indicates the fiber orientation with respect to the sample surface plane. The yellow square is the 1.5 × 1.5 mm^2^ region of interest for the orientation analysis.

**Fig 2:**
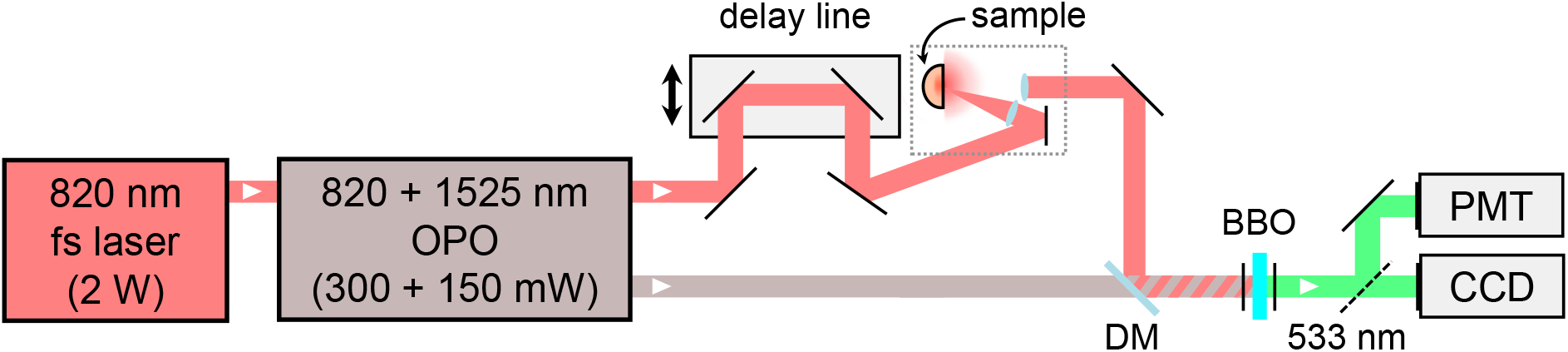
Sketch of the experimental setup for time- and space-resolved measurements when using a 820 nm probing wavelength. A Titanium-Sapphire fs laser is used to pump an optical parametric oscillator (OPO). The probe and gate beams are recombined using a dichroic mirror (DM) onto a *β*-barium borate (BBO) non-linear crystal, generating a sum-frequency signal that is detected with either a charge-coupled device (CCD) camera or a photomultiplier tube (PMT). The probe pulse impinges on the sample at an off-axis angle of *θ* = 13°, while light is collected perpendicularly to the sample surface, thus excluding light that is specularly reflected at the sample interface.

A series of 16 transient imaging frames were recorded with a CCD camera at different times between 0.5–15.5 ps with 1 ps incremental steps. The measurements were taken over a circular field of view with an area of 13.2 mm^2^, averaging over different adjacent positions across a ∼ 1 mm^2^ white matter region to reduce noise and average over speckle to obtain an incoherent profile. Within this limited time range, the sample can be effectively considered as homogeneous and semi-infinite since the observed diffused patterns remain bound within the white matter region both in the transverse plane and along the depth direction.

Notably, the measured profiles are independent of absorption, as the presence of absorption results in a mere modulation of their amplitude, leaving their functional shape unaffected. Quantitative information on the absorption coefficient can be retrieved by looking at the integrated time-resolved intensity, which can be obtained either by directly accumulating the intensity in each pixel, or using a photo-multiplier tube as detector for a more convenient sampling along the time axis.

### 3.3 Fiber orientation analysis

In order to correlate the anisotropic diffusion and scattering tensor components with the structural anisotropy, on the same sample, we perform an orientation analysis using a custom made light-sheet fluorescence microscopy (LSFM).^34^ In detail, with a vibratome (*Leica VT1000 S*) we cut a 1 mm-thick slice from the sample surface and we treat it with the *SHORT-DiD* protocol described by Sorelli et al.^35^ for myelinated fiber specific labeling and optical clearing of the tissue. The sample was imaged with the LSFM with an isotropic resolution of 3.6 µm and the 3D reconstruction was analyzed with the *Foa3D* tool^36^ producing a 3D fiber orientation map of the investigated white matter region with µm-scale resolution.

The analysis confirmed that the investigated white matter region is characterized by a homogeneous fiber density with a well-defined local alignment, allowing us to assume a spatially uniform scattering anisotropy tensor and reference frame for the diagonalization of the scattering and diffusion tensors, aligned along the local axon orientation. Fig. 1b shows the reconstructed fiber orientation in the sample surface plane. The system of reference for the analysis is then set having the *y* axis aligned with the average local orientation of the axon fibers in the area of interest, while the *x* axis is parallel to the sample surface. In this region, the fibers are found to have an elevation angle with respect to the sample surface of *α* = 0.5° with a standard deviation of *σ*_*α*_ = 8.7°. This means that the *xy* plane is basically aligned with the sample surface, which provides an ideal configuration to investigate the anisotropy. From now on, we will refer to the *y* axis as the parallel (∥) axis, and *x, z* as the perpendicular (⊥) axes to the axon fibers. Additionally, we will assume that the transport along the two perpendicular axes are the same, i.e., we will assume that the arrangement of axon fibers in the bundle is uniaxially symmetric on average around its main orientation axis.

## 4 Results

### 4.1 Transient imaging measurements

Experimental and simulated transient snapshots of the reflected intensity distributions are shown in Fig. 3. The profiles exhibit a faster diffusion rate along the direction parallel to the fibers, showing a clear hallmark of transport anisotropy. The profiles were fitted with a bi-variate Gaussian model to reconstruct the time evolution of their mean square displacement (MSD) growth along the parallel and perpendicular directions in the rotated system of reference (Fig. 4a). After the initial transient, both curves show a linear growth corresponding to the onset of diffusion, with different slopes associated to the different diffusion rates along the two axes. Performing a linear regression on the expansion rates for *t* > 10 ps, the components of the diffusive tensor are determined as: *D*_⊥_ = (1.69±0.06) × 10^−2^ mm^2^ ps^−1^ and *D*_∥_ = (2.12±0.11) × 10^−2^ mm^2^ ps^−1^, in perfect agreement with analogous measurements performed in transmittance on a slice of the same sample.^37^ Using these values with Eq. (4) allows us to evaluate the Optical Fractional Anisotropy of this white matter region, returning *OFA* = 0.135 ± 0.011. Notably, this value is lower than the water molecule diffusion *FA* usually found from DTI in white matter, which is around 0.4 in cortex white matter.^38, 39^

**Fig 3:**
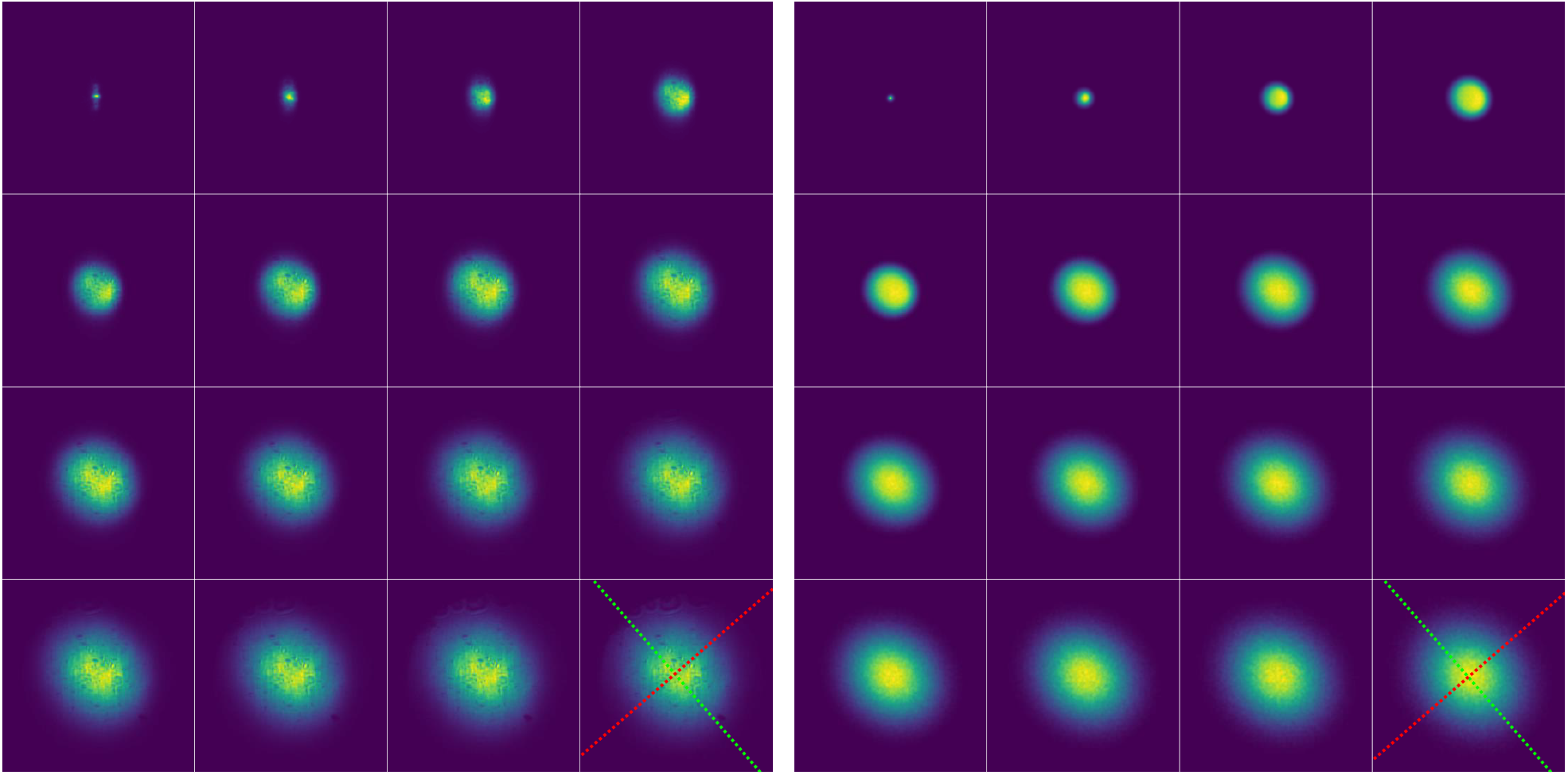
(a) Transient imaging measurements of the reflected intensity distribution, ordered from left to right, top to bottom, with increasing time delay step of 1 ps. Each frame is normalized to its maximum intensity and covers an area of 5.4 × 5.4 mm^2^. (b) Corresponding best-fit Monte Carlo simulation. The red and green dotted lines represent the axes perpendicular and parallel to the axon fibers, respectively.

**Fig 4:**
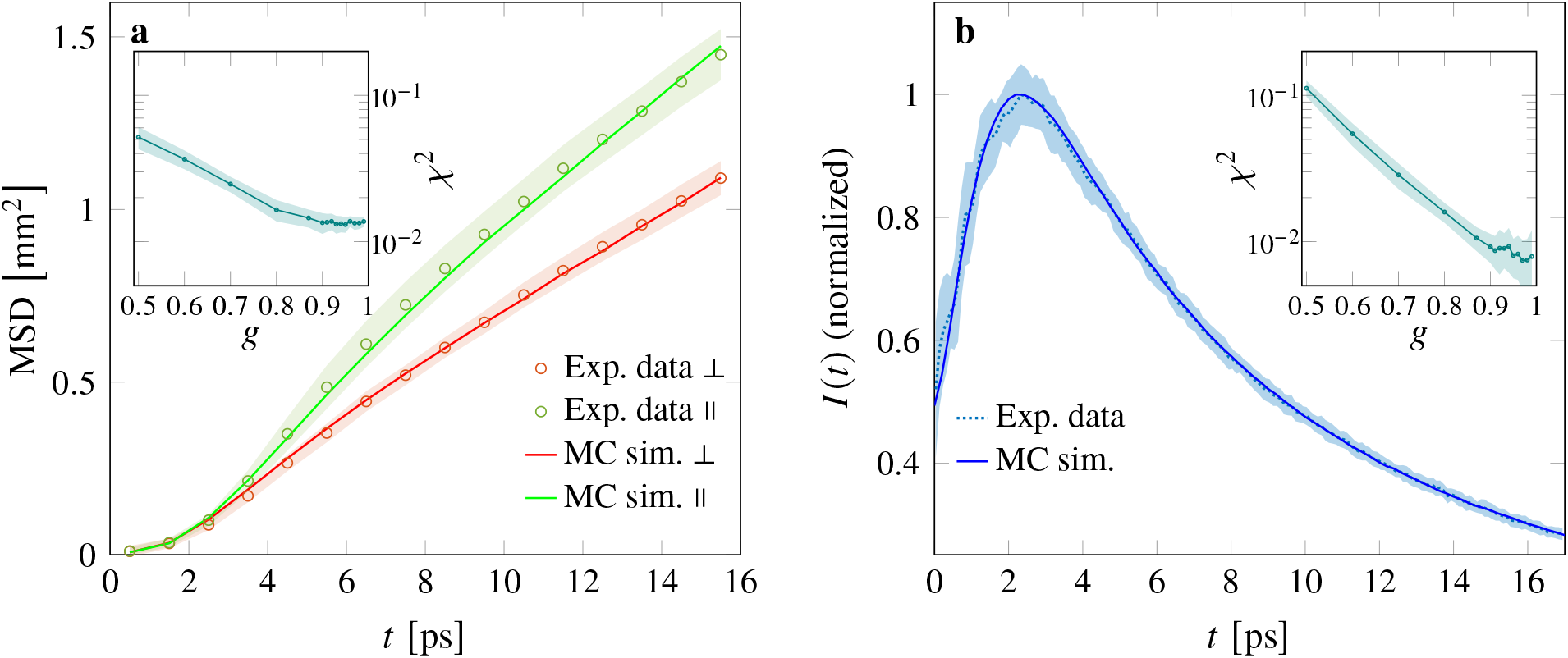
(a) MSD evolution along the perpendicular (⊥) and parallel (∥) directions. (b) Time-resolved reflected intensity. Solid lines represent the best-fit MC simulation. Shaded areas represent (a) the compounded uncertainty from the image acquisition and the bi-variate Gaussian fit, and (b) 1*σ* of the average between 5 different sample positions. The insets plot the fit *χ*^2^ dependence on *g* for the MSD evolution and time-resolved curves, respectively.

We fit the experimental data using simulations performed with *PyXOpto*, a general Monte Carlo software offering an implementation of the anisotropic tensor model.^40, 41^ The effective refractive index of white matter is assumed to be *n*_eff_ = 1.39, according to literature.^15, 42^ Taking advantage of the exact separation between scattering and absorption offered by the transient imaging data, the fitting procedure is performed in two stages. First, mean square displacement curves are analyzed using *μ*_s,∥_, *μ*_s,⊥_ and *g* as free parameters. Once these parameters are fixed, absorption is finally determined from the spatially-integrated time-domain reflectance curve, which also provides an independent confirmation of the fitted *g* value. The simulated results are shown in Figure 3 next to the experimental frames, exhibiting good agreement as confirmed by the comparison of the MSD curves (Fig. 4a). While the fit on the MSD linear region depends mostly on the effective reduced scattering coefficients, the early time transient (*t* < 8 ps) carries information on the single scattering properties. The goodness of the fit, expressed through its *χ*^2^ value, exhibits a clear dependence on the scattering asymmetry *g*, with best agreement obtained for *g* > 0.9 as expected or biological tissues. At such large asymmetry values, a small uncertainty on *g* can translate into a large variation of the retrieved scattering coefficients. In this regime, the effective reduced scattering coefficients represent a more robust set of parameters, returning 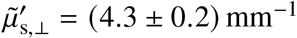 and 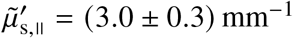. It is worth noting that the value of 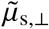, is consistent with the most comprehensive estimations of the (assumed isotropic) reduced scattering coefficient for white matter reported in the literature, ranging between 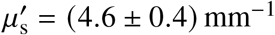 ^13^ and 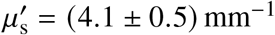 ^14^ at our test wavelength of *λ* = 820 nm. We ascribe this agreement to the fact that for a uniaxial anisotropic media, two out of three directions are characterized by the perpendicular scattering properties, which will therefore have a larger weight on the average transport properties. Nonetheless, our results show that scattering is markedly reduced along the prevalent alignment direction of fiber axon bundles, leading to an enhanced diffusion rate parallel to their orientation.

### 4.2 Time-resolved measurements

Given the retrieved components of the effective scattering mean free path tensor, we can analyze the integrated time-resolved data to determine the absorption coefficient and obtain an independent estimation of the asymmetry factor, which is known to influence the early-time transient of time-resolved reflectance curves.^43^ The fitting procedure (Fig. 4b) returns an absorption coefficient of *μ*_a_ = (0.022±0.008) mm^−1^. Previously reported values of the absorption coefficient for white matter are typically larger, in the range of *μ*_a_ = 0.07–0.09 mm^−1^,^13, 14^ which would cause time-domain curves to fall much more rapidly than what we observe experimentally. This discrepancy could be explained by the fixation of the sample, that has been shown to slightly reduce absorption,^16^ or by a slight overestimation of absorption in the previous literature. Regarding the scattering asymmetry, the minimum *χ*^2^ for this fit is again obtained for *g* = 0.9–0.99, with a minimum at *g* = 0.97, in excellent agreement with the independent MSD estimation. Notably, this result is also consistent with previous estimates in porcine and bovine brain,^11^ whereas it suggests that the widely assumed value for human brain white matter (*g* = 0.87)^13^ is underestimated. The high sensitivity of our time-resolved approach allows to discriminate between the single scattering effects at short times, and the effect of absorption at longer times, allowing for the simultaneous retrieval of these two parameters.

### 4.3 Estimation of the impact of white matter anisotropy in human head in fNIRS measurements

For practical functional near infrared spectroscopy application, it is interesting to evaluate the potential impact of ansisotropic diffusion in the white matter layer. In fact, when using non-invasive techniques, the effect of transport anisotropy in the white matter layer may be partially masked by the presence of the other intervening layers. To this purpose, we compare the results of two MC simulations in a representative four-layer model of the human head, assuming two orthogonal orientations of the white matter layer (Fig. 5a). The four layers represent respectively: the scalp/skull; the cerebrospinal fluid (CSF); gray matter (GM) and white matter (WM). For the first three layers, isotropic scattering and absorption properties are taken from the literature for a wavelength *λ* = 830 nm,^44^ while the WM layer is modeled using the experimentally determined parameters and anisotropy. The properties of all layers are summarized in Table 1.

**Table 1:** Properties of the four-layer used for the human head Monte Carlo simulation.^13, 44, 45^ For the white matter we use the experimentally determined values, replacing 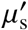 with the direction-dependent effective reduced scattering coefficients 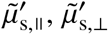.

| | Thickness [mm] | $n$ | $g$ | $\mu_a [\text{mm}^{-1}]$ | $\mu'_s [\text{mm}^{-1}]$ |
| --- | --- | --- | --- | --- | --- |
| Skull/Scalp | 10 | 1.39 | 0.89 | 0.019 | 0.86 |
| Cerebrospinal fluid | 2.5 | 1.34 | 0.89 | 0.0026 | 0.16 |
| Gray matter | 3 | 1.39 | 0.9 | 0.03 | 0.70 |
| White matter | 10 | 1.39 | 0.97 | 0.022 | 3.0/4.3 |

**Fig 5:**
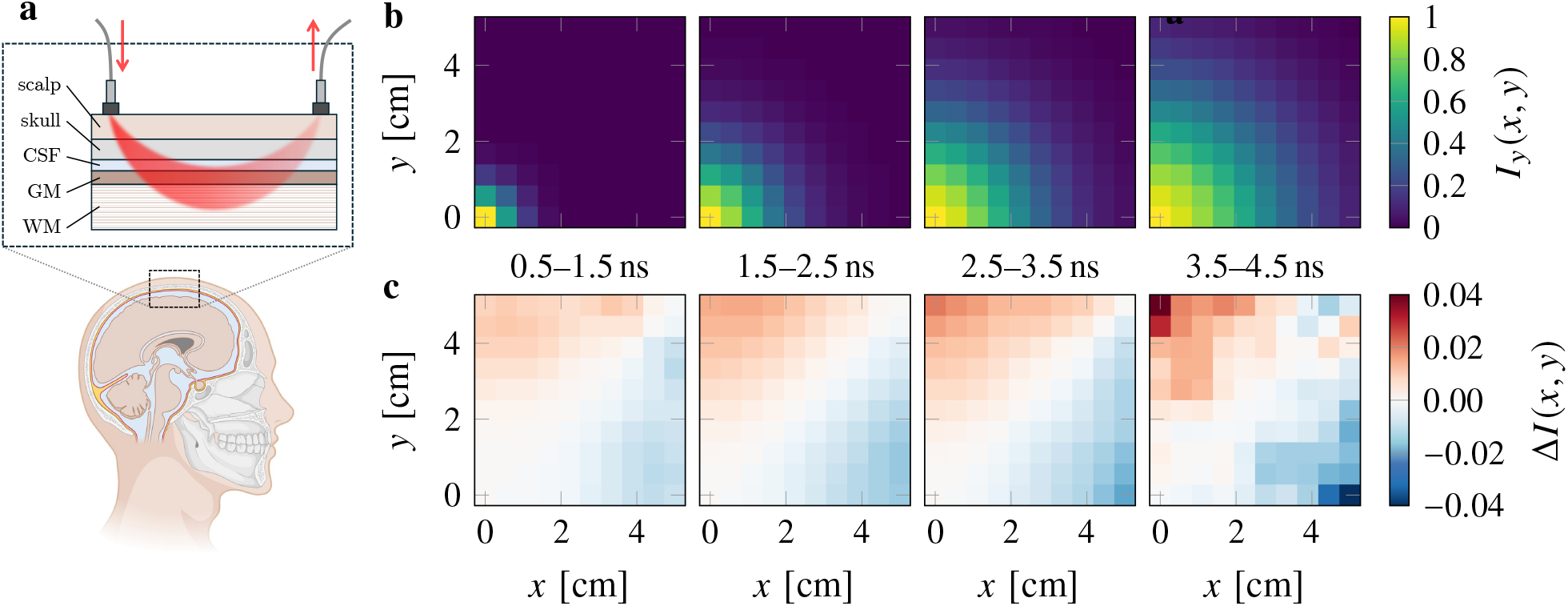
(a) Schematics of the four-layers model used in the MC simulation (created with BioRender). (b) Space-resolved reflectance at different time delays. Light is injected at position (0, 0) mm and collected over a 10 × 10 grid with 50 mm side (representing the upper right quadrant next to the illumination point). The faster diffusion axis of the underlying white matter layer is aligned with the *y* axis. (c) Relative difference between the space-resolved reflectance Δ*I* (*x, y*) of two simulations with orthogonal white matter anisotropy orientation.

In the simulation, the space- and time-resolved reflectance *I* (*x, y, t*) is collected from a 100 × 100 mm^2^ region around a central point illuminated with a delta-like pencil beam pulse. This configuration is analogous to a time-domain fNIRS measurement *in vivo* performed with optical fibers places at different positions from the illumination point, as schematically depicted in Fig. 5a. Two simulations are performed with the white matter fast-diffusion axis aligned with either the *y* and *x* axis, resulting in simulated intensities *I*_*y*_ (*x, y, t*) and *I*_*x*_ (*x, y, t*), respectively. The results of the two simulations, each comprising 10^11^ trajectories, are compared by taking their relative difference over time intervals at increasing delays between 0.5 ns to 4.5 ns, by calculating the quantity

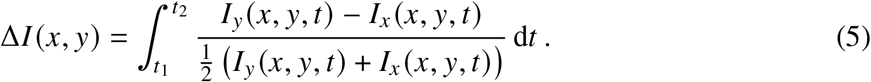

The result is shown in Fig. 5b, where we see that the overall difference increases with the distance and time delay from the illumination point. As expected, the effect of transport anisotropy in the deep WM layer is less pronounced when mediated through the outer head layers. Yet, the overall excess or depletion of light can reach a few percent points along the two directions, even when assuming that all the other layers are isotropic.

## 5 Discussion

This study presents the first quantitative evaluation of the anisotropic scattering tensor components of human brain white matter, linking it directly with the local axon orientation. Anisotropic MC simulations are shown to correctly reproduce both the experimental transient intensity profiles and time-resolved reflectance, including very early times associated to single scattering properties such as the asymmetry factor *g*. Our results show that the (isotropic) scattering coefficient values typically reported in the literature for human white matter are mostly influenced by the scattering coefficient perpendicular to axon fiber bundles, while a faster diffusive rate is present in the longitudinal direction.

These findings have implications for neuroimaging techniques such as functional near-infrared spectroscopy and diffuse optical tomography. Current models assume isotropic scattering, which may lead to inaccuracies in the interpretation of hemodynamic responses. By incorporating anisotropic scattering properties into these models, we can improve the accuracy of cerebral oxygenation measurements. This could enhance the diagnostic value of fNIRS and DOT, particularly in clinical applications where precise mapping of brain activity is critical.

In the field of optogenetics, accurate modeling of light transport is essential for effective neural stimulation.^8^ Our results suggest that accounting for anisotropic scattering could improve the design of optical stimulation protocols, ensuring that light reaches the intended target neurons with the desired intensity and spatial precision. This could enhance the efficiency of optogenetic activation, particularly for deep-brain stimulation, where light scattering significantly affects the illumination profile.

The same procedure applied here can be easily implemented to investigate light transport in other structurally anisotropic tissues, such as muscles^46, 47^ and tendons,^48, 49^ where directional dependence in light propagation is also typically disregarded when analyzing experimental data, even thought it could carry valuable information on the local degree of fiber organization allowing to discriminate between healthy and cancerous tissue.^50^

Also in the human head, additional anisotropy-induced effects may arise considering more realistic brain models, owing for instance to the fact that other brain tissues besides WM could be structurally anisotropic, or that the outer WM architecture exhibits U-shaped tracts that are often terminating with a perpendicular orientation to the skull. With enough sensitivity, these structures might provide a preferential path for probe light to reach a detector inadvertently positioned at the distal end of the tract. Other techniques with enhanced depth sensitivity such as the Dual-slope method^51, 52^ could also benefit from a correct accounting of the tensor scattering properties of WM tracts.

On top of this, deeper WM regions such as the corpus callosum or the spinal cord are also known to have an even more pronounced structural anisotropy in their axon fiber organization. We expect that these region would exhibit a higher *OFA*, leading to a correspondingly larger difference in the directional diffusion rates. Given the large extent of these highly aligned bundles connecting different regions throughout the brain, structural anisotropy may have a major impact on the probability of detecting diffuse photons even through the whole adult head.^53^ Similar effects could be measured also in infants due to their thinner skull allowing for deeper light penetration into the WM tracts and smaller head size,^54, 55^ even though in this case the *OFA* may be lower on average as axon fibers myelination progressively increases with age.^56^

In general, the existence of a direct correlation between the local axon bundle arrangement and the orientation of the co-located scattering tensor revealed the presence of a diffusive light transport enhancement along the main alignment direction of the white matter tracts. This result opens to the possibility of realizing a more accurate and anisotropy-aware 3D mesh of the human head for volumetric light scattering simulations, building on two-photon fluorescence microscopy 3D orientation analysis^36, 57^ or DTI tractography datasets^9, 58^ as a proxy to the local average orientation of fiber bundles at each location, which could help improving the accuracy of time-resolve transmittance measurements through the human head.^53, 59^

## Disclosures

The authors declare no competing interests.

## Code, Data, and Materials Availability

All the data that support the findings of this study are available from the corresponding authors upon request. Nerve fiber orientation analysis and anisotropic Monte Carlo simulations have been performed using the open-source software tools https://github.com/lens-biophotonics/Foa3D and https://github.com/xopto/pyxopto, respectively.

## Acknowledgments

This work was partially funded by the European European Union’s NextGenerationEU Programme with the I-PHOQS Research Infrastructure [IR0000016, ID D2B8D520, CUP B53C22001750006] “Integrated infrastructure initiative in Photonic and Quantum Sciences”. E.P. acknowledges financial support from Sony Europe B.V.. L.P. acknowledges the CINECA award under the ISCRA initiative, for the availability of high performance computing resources and support (ISCRA-C “ARTTESC2”). This project received funding from the General Hospital Corporation Center of the National Institutes of Health under award number U01 MH117023 and BRAIN CONNECTS (award number U01 NS132181). The content of this work is solely the responsibility of the authors and does not necessarily represent the official views of the National Institutes of Health-USA. From the European Union’s Horizon 2020 research and innovation Framework Programme under grant agreement No. 654148 (Laserlab-Europe), from HORIZON-INFRA-2022-SERV-B-01 “EBRAINS 2.0: A Research Infrastructure to Advance Neuroscience and Brain Health” Horizon Europe – Framework Programme for Research and Innovation (2021-2027). This research has also been supported by the Italian Ministry for University and Research in the framework of the Advanced Light Microscopy Italian Node of Euro-Bioimaging ERIC and by the European Union – Next Generation EU, Mission 4 Component 1, CUP B53C22001810006, Project IR0000023 SeeLife Strengthening the Italian Infrastructure of Euro-Bioimaging. The work was also supported by Fondazione Cassa di Risparmio di Firenze (project Human Brain Optical Mapping), by RICTD2025-2026 - CUP B97G24000240005, by LENS and CNR for the technical and scientific support to the Italian National Node FOE 2022 - CUP B53C24004790001. From University of Florence (D.R. n. 464 del 02/04/2024) with the project “Smart hydrogels with enhanced toughness to enable human brain tissue clearing (SMART-brain)”, CUP B97G24000240005.

The authors further thank Fabrizio Martelli and Alwin Kienle for fruitful discussion, and Christophe Destrieux from the Université de Tours for providing the human brain specimen analyzed in this study. We express our gratitude to the donor involved in the body donation program of the Association des dons du corps du Centre Ouest, Tours, who made this study possible by generously donating his body to science.

**Ernesto Pini** is a PostDoc at the Italian National Institute for Metrological Research since 2024. His PhD was funded by SONY Europe B.V. and it was conducted in collaboration with their research center in Stuttgart. He received his BS and MS degrees in physics from the University of Florence in 2018 and 2021, respectively. His current research concerns light transport in disordered materials, encompassing interests in Photonic materials, Biophotonics and Transport theory. He is a member of SPIE.

## Notes

### Competing Interest Statement

The authors have declared no competing interest.

